# Magnetic fluctuations affect circadian patterns of locomotor activity in zebrafish

**DOI:** 10.1101/2021.09.08.459369

**Authors:** Viacheslav V. Krylov, Evgeny I. Izvekov, Vera V. Pavlova, Natalia A. Pankova, Elena A. Osipova

## Abstract

The locomotor activity of zebrafish (*Danio rerio*) has a pronounced, well-studied circadian rhythm. Under constant illumination, the period of free-running locomotor activity in this species usually becomes less than 24 hours. To evaluate the entraining capabilities of slow magnetic variations, zebrafish locomotor activity was evaluated at constant illumination and fluctuating magnetic field with a period of 26.8 hours. Lomb-Scargle periodogram revealed significant free-running rhythms of locomotor activity and related behavioral endpoints with a period close to 27 hours. Obtained results reveal the potential of slow magnetic fluctuations for entrainment of the circadian rhythms in zebrafish. The putative mechanisms responsible for the entrainment are discussed, including the possible role of cryptochromes.

## INTRODUCTION

Circadian rhythms play a significant role in the physiology of the majority of living beings. They provide effective use of energy and resources in ever-changing natural and artificial environments [1]. Based on the endogenous rhythms of intracellular circadian oscillators, an organism adjusts its internal processes to the anticipated conditions for a given time of day [2]. Briefly, these circadian clocks in cells are described as transcription-translation feedback loops. In most vertebrates, positive components of this loop are the transcription factors CLOCK and BMAL that modulate the expression of Period (*Per*) and Cryptochrome (*Cry*) genes as negative components. These negative components repress transcription and induce the body’s circadian clock to reset, thus starting a new cycle of the feedback loop [1, 3, 4]. In birds and mammals, this endogenous circadian oscillator (located in the brain’s suprachiasmatic nucleus) provides the main rhythm transferred to peripheral tissues via pineal gland produced melatonin [5]. A decentralized to varying degree circadian system can be found throughout the evolutionary tree [6, 7].

The endogenous circadian rhythms adjust to external environmental cues (zeitgebers) with the primary external pacemaker being light/dark cycles. In general terms, the endogenous circadian oscillator synchronizes to local daytime via photic cues transmitted from the retina to neurons of the suprachiasmatic nucleus [8]. Some findings suggest the diurnal geomagnetic variation may be a secondary external zeitgeber affecting biological circadian rhythms [9, 10]. This variation results from the dynamo-current process within the ionospheric E-region and represents a distinguishable daily magnetic oscillation from approximately a few tens of nanoTeslas (nT) at mid-latitudes to 200 nT near the magnetic equator [11]. Diurnal geomagnetic variation is suggested to act as a potential circadian zeitgeber via cytochromes possibly being able to perceive magnetic fields through radical pair reactions [12, 13]. Though indirect evidence supports this theory [14, 15, 16, 17, 18, 19, 20], no direct experimentation has been carried out studying the entrainment of circadian rhythms to slow magnetic fluctuations. The issue of whether slow changes in the magnetic field affect circadian oscillators remains open.

In the present study, we used wild-type zebrafish (*Danio rerio*) to answer this question. The brain of this species contains the pineal gland driving the rhythmic production of melatonin [21, 22]. However, the circadian oscillators in different zebrafish tissues can keep unrelated rhythms that are entrained directly by an external light-dark zeitgeber [23, 24, 25]. The lack of a centralized pacemaker subjugating all other oscillators in zebrafish could increase the chance to detect changes in circadian rhythms caused by a magnetic influence. The influence of different light and magnetic exposures on zebrafish rhythms was analyzed through locomotor activity and several related endpoints known as precise indicators of circadian rhythmicity in this species [26].

## METHODS

All animal experiments were carried out in accordance with relevant guidelines and regulations. The study was carried out in compliance with the ARRIVE guidelines. All experimental protocols have been approved by the Institutional Animal Care and Use Committee at Papanin Institute for Biology of Inland Waters (https://ibiw.ru/index.php?p=downloads&id=46180).

### Zebrafish maintenance

Wild-type zebrafish (AB strain) were obtained from the commercial distributer and maintained in the Laboratory of physiology and toxicology (Papanin Institute for Biology of Inland Waters, Russian Academy of Sciences). Prior to experimentation, zebrafish were kept together for two months in 70 L aquaria at 24°C under a 16:8 h light/dark cycle. Zebrafish were fed daily at different times between 12:00 and 16:30. Males and females at the age of approximately four months (mean body length 2.99 cm, SD = 0.17 cm, n = 24) were used for experimentation.

### Timed backlight

In order to provide backlit illumination for the experiments, a lightbox was constructed from a series of LEDs, aluminum plates, and matte plexiglass. LED plates were created by adhering 32 LEDs to an aluminum plate so as each aquarium would be backlit by 4 infrared LEDs (3W, 940nm) and 4 white-color LEDs (3W, 4500K). Each LED plate was mounted 10 cm under a lightbox cover (constructed from matte plexiglass) which serves to diffuse light. Lighting modes were controlled via time relays (DH-48S-S, Omron, Japan) which used KMI-10910 (IEK, Russia) contactors to supply power, by Qh-60LP18 power suppliers (Shenzhen Chanzon Technology, China), to the LEDs.

### Magnetic fluctuations

#### Zebrafish were exposed to the following magnetic fluctuations

##### 1. The natural diurnal geomagnetic variation

It is represented by magnetic fluctuations of about 30 nT with a 24 h period. This variation was recorded in X-, Y-, and Z-directions throughout the experiment using an NV0302A magnetometer (ENT, St Petersburg, Russia). Six geomagnetic disturbances with a k-index of 4 that corresponds to weak geomagnetic storms occurred during the experiments under natural diurnal geomagnetic variation (08/02/2020 from 12:00 to 18:00; 08/31/2020 from 00:00 to 03:00, from 09:00 to 12:00, and from 15:00 to 21:00; 09/14/2020 from 00:00 to 03:00). The natural diurnal geomagnetic variation was accompanied by a 16: 8 h light / dark photoperiod.

##### 2. Experimental magnetic fluctuations simulating increased diurnal geomagnetic variation with an average period of 26.8 h

We used a sample record of diurnal geomagnetic variation in X-, Y- and Z-directions made close to the laboratory to generate these magnetic oscillations. The sample record intensity was enhanced to about 100-150 nT for each X-, Y-, and Z-directions. This exposure allows for more pronounced periodic changes in the magnetic background but not exceeding the level of natural geomagnetic storms. The period of sample diurnal geomagnetic variation was also increased to 26.8 h by a signal prolongation. This value was chosen for the experiments because the free-running rhythm of locomotor activity in zebrafish became shorter than 24 h under constant illumination. Hence the magnetic zeitgeber with a period longer than 26 h can manifest itself under constant illumination. At the same time, the period of 26.8 h is quite close to the circadian 24 h period. That is, the endogenous oscillator does not require drastic changes in order to be entrained by this external zeitgeber. The experimental magnetic fluctuations were generated under constant light conditions. Signals of the natural diurnal geomagnetic variation and experimental magnetic fluctuations in the horizontal direction are shown in Figure 1. In order to compare these signals with behavioral endpoints, they are also presented as actograms and periodograms (Fig. 2 a, b).

**Fig 1.**
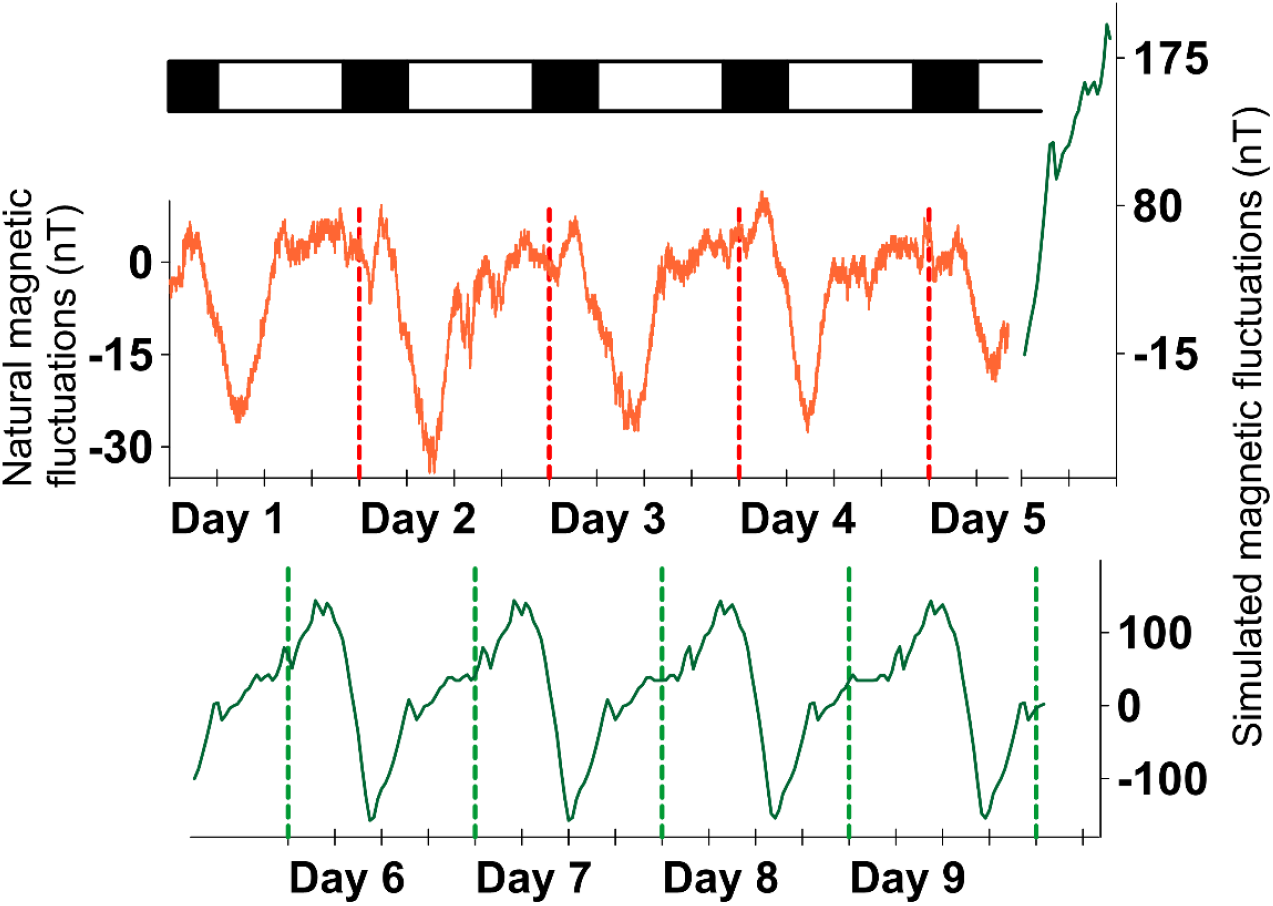
Natural diurnal geomagnetic variation (red line) and experimental magnetic fluctuations (green line) in the horizontal direction.

**Fig 2.**
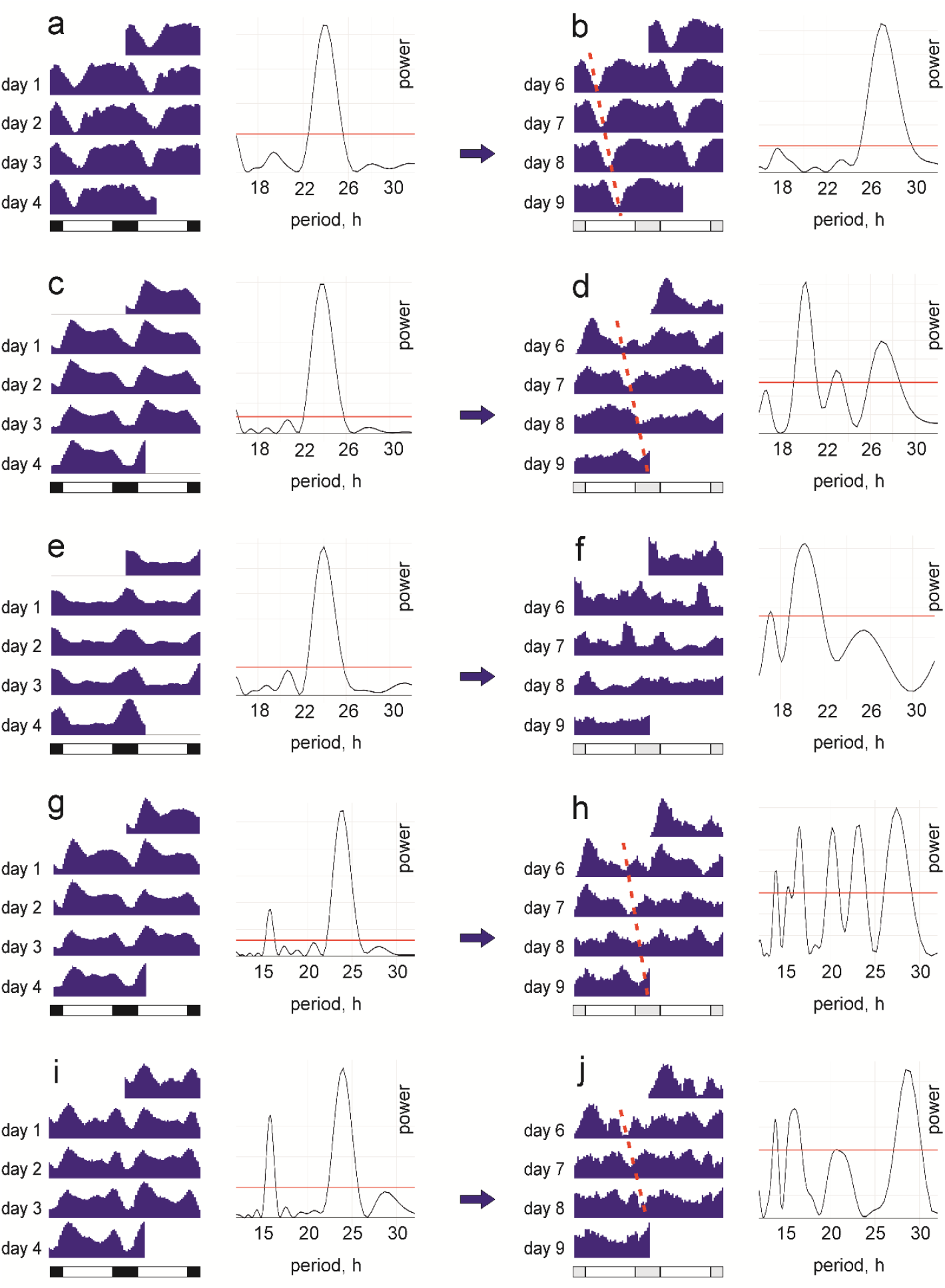
Diurnal geomagnetic variation (a), experimental magnetic fluctuations (b), and dynamics of zebrafish behavioral endpoints (c–j) given as a set of double plotted actograms and periodograms. Both natural geomagnetic variation used at the first stage of experiments under a 16:8 LD cycle (a) and experimental magnetic fluctuations with a 26.8 h period under constant illumination at the second stage (b) were horizontally directed. Behavioral endpoints measured at these two stages respectively included: average swimming speed (c, d), meandering (e, f), angular velocity (g, h), and wall preference index (i, j). Significant periods on Lomb-Scargle periodograms (p < 0.05) are above the solid red line. The dotted line on the actograms denotes the trends that determine a significant free-running period of about 27 hours.

A setup, described in detail by Krylov et al. [27], was used to generate experimental magnetic fluctuations. It was assembled on a PC workstation and consisted of the following items:

1. A three-component fluxgate magnetometer NV0302A (ENT, St Petersburg, Russia providing analogous signals proportional to the strength of the geomagnetic field and its variations;
2. An LTR11 analog-to-digital and an LTR34-4 digital-to-analog signal converter (L-card, Moscow, Russia);
3. A coil system consisting of three pairs of mutually orthogonal Helmholtz coils (0.5 m in diameter, 700 turns of 0.2 mm copper wire in each coil) made by the Schmidt Institute of Physics of the Earth (www.ifz.ru).

The direction of each Helmholtz coils pair was the same as the direction of the geomagnetic field components. Natural fluctuations of the geomagnetic field, including diurnal variation, were compensated within the Helmholtz coil systems in the frequency range up to 5 Hz based on a signal from NV0302A magnetometer (ENT). The industrial alternating magnetic fields of 50 Hz were less than 10 nT and did not appear in the harmonics. Parameters of the generated signals in the Helmholtz coils system’s working volume were checked using a control magnetometer NV0599C (ENT).

### Experimental conditions and procedure

All experimentation was conducted in a remote laboratory free of working staff in order to eliminate possible circadian rhythm influences caused by daily human activities. Four fish were placed in four custom glass aquaria (15 x 20 cm, height 23 cm) filled with 10 cm of water, with one fish per aquarium. Water temperature during the experiments was 21^°^C as adult zebrafish show the most robust rhythm of locomotor activity at the temperatures of 20-21 ^°^C (Hurd et al., 1998; López-Olmeda et al., 2006). The aquaria were installed above a lightbox. Screens made of opaque white plastic were placed between the adjacent aquaria so that fish could not see conspecifics. The lightbox with the aquaria was located in a system of Helmholtz coils. During the first 4.5 days, a 16: 8 h light / dark cycle was maintained, and no voltage was supplied to Helmholtz coils. Then from 13:00 of the 5th to 00:00 of the 10th day of the experiments, constant lighting conditions were maintained, and experimental magnetic fluctuations were generated within Helmholtz coils.

The water was constantly renewed via two 4 mm openings in the wall of each aquarium at 3 and 10 cm height from the bottom. Water flowed by gravity from a 200 L plastic barrel placed one floor above through the silicone hoses connected to the bottom openings of aquaria. Water aeration and temperature control for all aquaria were carried out in the barrel. Excess water was drained to the sewer through the top opening to ensure a constant level of 10 cm. Water from different aquaria has never been mixed or reused.

At the beginning of the experiment, a 1 cm^3^ piece of slow-release gel food block “Tetra Holiday” (Tetra GmbH, Melle, Germany) was placed on the bottom of each aquarium to prevent the influence of the feeding schedule on circadian behavior. Thereby zebrafish had free access to food during the whole study.

Fish movements in the horizontal plane were registered with IP-cameras (TR-D1140, Trassir, Shenzhen, China) equipped with IR corrected varifocal lenses (TR-L4M2.7D2.7-13.5IR, Trassir, Shenzhen, China) and mounted above the aquaria. Night and day video was recorded in black and white at 25 frames per second with a resolution of 2592 × 1520 pixels. The video signals were transmitted through a switch (T1500-28PCT, TP-Link, Shenzhen, China) to a video recorder server (MiniNVR AF16, Trassir, Shenzhen, China).

The experiment was performed in 3 independent and time-separated replications between July 31, 2020 and September 17, 2020. Each zebrafish was used only for a single replication. Thereby 216-h video records obtained from 12 zebrafish were then processed.

### Data processing

An approach proposed by Audira et al. [28] was used for data processing. One-minute video files were cut from the primary video record for every half of an hour (from the 15th to the 16th and 45th to 46th min of each hour). Such duration has proved to be sufficient for statistical analysis of locomotor activity with the data appropriately describing circadian rhythms [28, 29, 30]. The open-source software idTracker [31] was used to process each one-minute video file. The software provided X and Y coordinates reflecting the center of the fish body for each frame. Before the processing, the trajectory data were filtered using the “*minimal distance moved”* method to eliminate slight “apparent” movements of the fish [32, 33]. The minimal distance threshold was set at 2.6 mm. Then, based on this information, several quantitative measures of fish behavior that reveal pronounced circadian rhythms in zebrafish [28] were calculated using the Microsoft Excel formulae. The parameters included:

1. Average swimming speed, cm/s (total distance travelled divided by total observation time)
2. Meandering, °/cm, reflecting the trajectories irregularity and calculated as the sum of all turning angles (absolute values) divided by total distance
3. Average angular velocity, °/s (total turning angle divided by total test time)
4. Freezing time, % (the total time when speed is less than 1 cm/s)
5. Swimming time, % (the total time when speed ranges from 1 to 10 cm/s)
6. Rapid movement time, % (the total time when speed exceeds 10 cm/s)
7. Wall preference index (relative time spent within a 3-cm-wide area close to the walls)

The latter index is calculated upon the formula: 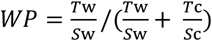, where *T*w and *T*c denote time (s) spent close to the walls and in the central zone, respectively, while *S*w and *S*c denote the area (cm^2^) of each zone. The index varies from 0 (if a fish never approaches the walls) to 1 (if a fish keeps the walls and never visits the central zone).

Differences between the average values of studied parameters during the light and dark phases were evaluated with a t-test as all data had a normal distribution (Shapiro–Wilk W-test, p > 0.05). Time series were analyzed with RhythmicAlly software [34]. The linear trend was subtracted from the time series, and the data were smoothed with a moving average window of 7 samples before analysis. Circadian periods under 16: 8 h light / dark cycle and free-running periods under constant illumination were analyzed using the Lomb-Scargle periodogram [35]. We also used cosinor-analysis [36] based on the approximation of a time series by a cosine wave to identify a mesor (or a rhythm-adjusted mean that represents the average level of the cosine wave) and an amplitude (a measure of half the extent of predictable variation within a cycle) of studied rhythms.

## RESULTS

Zebrafish displayed a robust circadian rhythm of locomotor activity and related behavioral endpoints at the first stage of the experiments under 16:8 h light / dark cycle. Most of the endpoints (swimming speed, angular velocity, wall preference index, swimming time, and rapid movement time) were higher during the light phase and lower in the dark, while meandering and freezing time followed the reversed pattern (Suppl. 1). The circadian period in the dynamics of studied endpoints was 24 h (Fig. 2 c, e, g, i). The average angular velocity and wall preference index had an additional weaker rhythm with a 15.84 h period (Fig. 2 g, i). Natural geomagnetic disturbances with a k-index of 4 did not affect circadian patterns of behavioral endpoints at the first stage of the experiments.

At the second stage of the experiments, the zebrafish were held at constant lighting and magnetic variation with a 26.8-h period (Fig 2 b). In the absence of the photic zeitgeber, the studied endpoints, except for meandering, showed significant free-running rhythms with periods close to 27 h (Fig. 2 d, f, h, j). These free-running rhythms were dominant for angular velocity, wall preference index, and rapid movement time (Table 1). However, in the case of swimming speed, freezing time, and swimming time, these 27 h rhythms, while present, were less pronounced than those with 20 h periodicity (Table 1). Generally, the amplitudes of behavioral rhythms found within the second stage of the experiments were reduced compared to the first stage through the emergence of additional rhythms manifested in several peaks on periodograms. Such a multiple-peaked pattern results from different individuals with a predominance of one or another rhythm in the studied group (Suppl. 2). However, even in individuals with a predominance of one of the most pronounced periods (20 ± 1 h or 27 ± 1 h), a secondary, less prominent peak was often present. In two individuals, these two peaks had almost equal amplitude. At the same time, several individuals in the group retained a rhythm with a period close to 24 h (Suppl. 2).

**Table 1.** Lomb-Scargle periods, mesors, and amplitudes for the rhythms of studied behavioral endpoints in zebrafish. All presented rhythms are significant (p <0.05, Lomb-Scargle periodogram analysis).

| Parameter | First stage of experiments,<br>circadian period |  |  | Second stage of experiments,<br>primary free-running period |  |  | Second stage of experiments,<br>secondary free-running period |  |  |
| --- | --- | --- | --- | --- | --- | --- | --- | --- | --- |
|  | period | mesor | amplitude | period | mesor | amplitude | period | mesor | amplitude |
| Average swimming speed | 24 | 1.852 | 0.635 | 20.211 | 1.998 | 0.103 | 26.947 | 2.020 | 0.100 |
| Meandering | 24 | 46.781 | 19.021 | 20.211 | 33.405 | 1.245 | 17.067 | 33.441 | 1.021 |
| Average angular velocity | 24 | 67.711 | 12.045 | 27.429 | 62.181 | 2.417 | 20.211 | 61.794 | 3.309 |
| Wall preference index | 24 | 0.494 | 0.030 | 28.444 | 0.571 | 0.011 | 16.000 | 0.570 | 0.010 |
| Freezing time | 24 | 41.746 | 23.703 | 20.211 | 25.526 | 1.566 | 26.947 | 25.119 | 1.673 |
| Swimming time | 24 | 57.445 | 23.529 | 19.948 | 73.985 | 1.488 | 26.947 | 74.390 | 1.610 |
| Rapid movement time | 24 | 0.809 | 0.625 | 27.927 | 0.515 | 0.094 | 20.480 | 0.496 | 0.101 |

The diurnal changes in studied behavioral endpoints significantly correlated with the experimentally generated magnetic fluctuations and were not related to the natural geomagnetic variation except for a couple of weak correlations (Table 2).

**Table 2.** Spearman rank-order correlations between the behavioral endpoints in zebrafish and magnetic fluctuations in X, Y, and Z directions during the second stage of experiments (n = 3348 for each correlation). The correlation coefficients between the behavioral endpoints and natural diurnal geomagnetic variation are also given for comparison. This geomagnetic variation would have been if the magnetic field had not been modified in the experiment. Significant correlations at p < 0.001 (after correction for multiple pairwise correlations) are marked with asterisks.

| Parameter | Experimental magnetic fluctuations |  |  | Natural diurnal geomagnetic variation |  |  |
| --- | --- | --- | --- | --- | --- | --- |
|  | X | Y | Z | X | Y | Z |
| Average swimming speed | -0.204* | 0.116* | -0.114* | -0.064 | -0.113* | -0.047 |
| Meandering | 0.182* | 0.333* | -0.160* | -0.027 | 0.028 | 0.033 |
| Average angular velocity | -0.031 | -0.582* | 0.386* | -0.026 | -0.043 | -0.043 |
| Wall preference index | -0.235* | 0.415* | -0.355* | -0.049 | -0.040 | -0.014 |
| Freezing time | 0.182* | -0.207* | 0.191* | 0.033 | 0.046 | 0.000 |
| Swimming time | -0.178* | 0.219* | -0.195* | -0.029 | -0.046 | -0.002 |
| Rapid movement time | -0.107* | -0.049 | 0.050 | -0.066 | -0.111* | -0.064 |

## DISCUSSION

The circadian rhythms in the dynamics of the studied parameters at the first stage of the experiment correspond to the known patterns of zebrafish circadian behavior governed by daily changes in illumination [26]. Additional peaks at 15.84 h on the periodograms for the wall preference index and angular velocity are associated with a well-documented phenomenon of visual-motor response in zebrafish [37]. In the present case, fish preferred walls to the inner area of the aquarium and showed increased angular velocity at the moments of abrupt changes in illumination.

It is known that the period of free-running rhythms in zebrafish locomotor activity under constant illumination and not-modified geomagnetic conditions usually becomes shorter than 24 h. Thus, Hurd et al. [38] reported that such free-running periods vary in the range of 23.5-24.5 h depending on the water temperature. Another study revealed the shortening of daily rhythms in zebrafish locomotor activity to 22.9-23.6 h under constant dim light [39]. The free-running rhythm of locomotor activity in zebrafish also became shorter (22.9 ± 0.5 h) under ultradian 45:45 min light/dark cycles [40]. We found no mention of the zebrafish locomotor activity rhythms with a period longer than 26 h maintained under constant illumination without additional zeitgebers. Significant 27 h peaks found on the fish periodograms in the present study coincide with the period of experimental magnetic fluctuations. These data strongly suggest that the slow changes in the external magnetic field may entrain the free-running behavioral rhythms in zebrafish. This is also evidenced by significant correlations between the studied here behavioral endpoints and experimental magnetic fluctuations. Earlier, it was reported that magnetic influence could affect circadian rhythms in different organisms [41, 42]. However, until the present study, there were no direct experimental data in support of the entrainment of endogenous circadian oscillators to slow magnetic fluctuations.

At the same time, zebrafish showed pronounced individuality of the entrainment. These results are in accordance with previously reported data on the variability of zebrafish behavioral responses in general [38, 43] and marked individuality of their reactions to magnetic fields in particular [44, 45].

Our results also indicate that under constant illumination in the presence of a 26.8-hour magnetic zeitgeber, competition likely occurs between the two free-running rhythms found in zebrafish. One of these rhythms follows the magnetic zeitgeber and has a visible period of about 27-h. Apparently, cryptochromes could participate in the entrainment of locomotor activity rhythms with magnetic fluctuations. On the one hand, it was suggested that cryptochromes are responsible for the biological effects of geomagnetic storms [10] and magnetic-compass orientation [13]. On the other hand, cryptochromes are involved in the transcription-translation feedback loop as the main elements of the molecular circadian oscillator [1, 46]. Some investigations revealed a direct link between the magnetic field intensity and expression levels of cryptochromes [16] and other clock genes [18]. A possible mechanism of magnetic influence on cryptochromes is based on changes of the singlet-triplet state of electrons in cryptochrome’s radical pairs, modulating the functional state of these proteins [12]. These magnetic-field-induced changes in the functional state of cryptochromes may, in turn, affect the repressor functions of the CRY:PER dimers. At the same time, other elements of the complex molecular circadian oscillator network [46, 47] may continue to function in a usual mode, which would shorten the free-running period under constant illumination. Due to these processes, two or more free-running rhythms with different periods can arise under the experimental magnetic influence. In addition, zebrafish possess various independent cellular oscillators with diverse rhythms in different tissues [7, 23, 48]. It can also be the reason for several periodogram peaks found in the present experiments.

Circadian patterns of behavioral endpoints at the first stage of the experiments depended on the light-dark cycle. They were not affected by natural disruptions of diurnal geomagnetic variation with a k-index of 4. The magnetic zeitgeber manifested itself only in the absence of a light-dark cycle in the second stage of the experiment. Hence changes in illumination have a greater impact on circadian patterns of zebrafish locomotor activity than magnetic fluctuations.

It needs to be emphasized that the present results are significant for the time series obtained in this experiment. Further research needs to be performed considering the individuality in zebrafish responses to magnetic influence under constant illumination. The present results indicate a high possibility of the entrainment of circadian rhythms to slow magnetic fluctuations. More experiments from different scientific groups are needed to clarify this issue and expand our knowledge of non-photic cues for periodic biological processes. It can opens prospects for manipulating circadian oscillators via magnetic fields. Further research in the field can focus on studying the effects of slow magnetic fluctuations on circadian genes’ rhythmic expression. Some recommendations for generation of slow magnetic fluctuations are given in Supplementary 3.

The datasets analyzed during the current study are available in the “Open Science Framework” repository, https://osf.io/4by9t/

## ACKNOWLEDGMENTS

The authors express their sincere gratitude to G.M. Chuiko for the provided wild-type zebrafish. We thank O.D. Zotov and B.I. Kline for help in preparing experimental signals and processing time-series data, M.A. Tyumin for technical support, and O.V. Solovieva for her help in the data processing. This research received support from the Russian Foundation for Basic Research (project No. 20-04-00175) to V.V.K., E.I.I., V.V.P., and N.A.P. E.A.O. was supported by the Ministry of Education and Science of the Russian Federation (State Assignment No. 121051100109-1).

## AUTHOR CONTRIBUTIONS

V.K. conceived and planned the experiments. V.K. and E.I. carried out the experiments. All authors processed the experimental data and performed the analysis. V.K. and E.I. wrote the paper with input from all authors.

