## Supplementary material for "Magnetic fluctuations affect circadian patterns of locomotor activity in zebrafish": Suppl

**Supplementary 1**

**Table S1.** Studied behavioral endpoints in zebrafish at the light (over the dash) and dark (under the dash) phases during the first stage of experiments

| Individual | Average swimming speed (cm/s) | Meandering (°/cm) | Average angular velocity (°/s) | Wall preference index | Freezing time (%) | Swimming time (%) | Rapid movement time (%) | n |
|---|---|---|---|---|---|---|---|---|
| 1 | 2.41±0.09<br>1.44±0.16 * | 34.64±1.26<br>76.60±7.09 * | 86.86±5.50<br>70.28±6.83 | 0.54±0.02<br>0.38±0.04 * | 16.07±1.60<br>62.78±4.22 * | 83.03±1.60<br>36.35±4.12 * | 0.91±0.15<br>0.87±0.15 | 134<br>76 |
| 2 | 2.35±0.09<br>1.04±0.12 * | 31.07±1.40<br>74.41±6.04 * | 70.63±4.01<br>53.86±5.59 * | 0.43±0.02<br>0.29±0.04 * | 26.54±2.01<br>69.84±3.82 * | 72.91±1.98<br>29.90±3.79 * | 0.55±0.09<br>0.26±0.05 * | 135<br>76 |
| 3 | 3.68±0.16<br>1.54±0.15 * | 32.91±1.78<br>49.91±2.46 * | 117.07±6.46<br>64.21±5.00 * | 0.58±0.03<br>0.55±0.04 | 8.06±1.20<br>57.23±3.54 * | 86.81±1.29<br>42.25±3.46 * | 5.13±0.83<br>0.52±0.12 * | 135<br>76 |
| 4 | 3.25±0.10<br>1.40±0.14 * | 25.37±0.88<br>57.89±4.96 * | 84.39±4.22<br>62.63±6.43 * | 0.61±0.02<br>0.41±0.05 * | 9.22±0.98<br>59.20±4.04 * | 89.19±0.96<br>40.49±4.00 * | 1.58±0.23<br>0.30±0.06 * | 135<br>76 |
| 5 | 1.36±0.05<br>0.84±0.08 * | 38.97±2.43<br>108.66±11.37* | 46.32±2.35<br>52.07±4.09 | 0.42±0.02<br>0.64±0.04 * | 43.54±2.32<br>69.80±3.30 * | 56.40±2.32<br>30.13±3.29 * | 0.06±0.01<br>0.07±0.02 | 133<br>76 |
| 6 | 2.32±0.11<br>0.77±0.06 * | 38.42±1.38<br>49.64±2.84 * | 84.88±4.78<br>31.22±1.40 * | 0.49±0.03<br>0.29±0.03 * | 29.92±2.65<br>76.00±2.28 * | 68.98±2.60<br>23.88±2.26 * | 1.10±0.20<br>0.12±0.02 * | 134<br>76 |
| 7 | 1.34±0.07<br>0.67±0.06 * | 32.68±1.76<br>66.67±7.82 * | 39.87±2.40<br>25.49±2.10 * | 0.33±0.02<br>0.57±0.04 * | 52.00±2.53<br>79.26±2.35 * | 47.85±2.53<br>20.63±2.34 * | 0.14±0.03<br>0.11±0.02 | 133<br>76 |
| 8 | 2.09±0.09<br>0.87±0.09 * | 35.84±0.89<br>76.64±5.41 * | 76.46±4.17<br>46.23±3.79 * | 0.43±0.03<br>0.39±0.05 | 30.80±2.21<br>72.55±3.44 * | 68.84±2.20<br>27.22±3.43 * | 0.36±0.06<br>0.23±0.06 | 134<br>76 |
| 9 | 3.67±0.10<br>0.91±0.04 * | 45.51±1.00<br>65.60±3.55 * | 162.39±4.78<br>56.43±2.98 * | 0.83±0.01<br>0.57±0.03 * | 10.87±0.76<br>70.84±1.82 * | 86.13±0.67<br>29.03±1.82 * | 3.00±0.35<br>0.13±0.03 * | 135<br>76 |
| 10 | 1.94±0.07<br>0.70±0.06 * | 35.83±1.42<br>68.71±5.52 * | 65.58±3.08<br>35.64±2.61 * | 0.53±0.02<br>0.51±0.04 | 26.13±1.94<br>79.50±2.14 * | 73.50±1.93<br>20.34±2.12 * | 0.37±0.09<br>0.16±0.04 | 135<br>76 |
| 11 | 2.06±0.05<br>0.87±0.11 * | 27.48±0.71<br>52.63±5.38 * | 56.33±1.81<br>36.60±3.53 * | 0.56±0.02<br>0.36±0.03 * | 16.73±0.90<br>75.10±3.12 * | 83.03±0.90<br>24.76±3.10 * | 0.24±0.03<br>0.13±0.04 * | 135<br>76 |
| 12 | 1.45±0.08<br>0.50±0.04 * | 45.06±2.93<br>89.85±7.28 * | 52.40±2.53<br>38.01±3.18 * | 0.48±0.02<br>0.47±0.04 | 46.99±2.39<br>85.42±1.74 * | 52.64±2.36<br>14.51±1.73 * | 0.37±0.15<br>0.06±0.01 | 135<br>76 |
| average (n=12) | 2.33±0.03<br>0.96±0.03 * | 35.31±0.49<br>69.77±1.88 * | 78.68±1.42<br>47.56±1.31 * | 0.52±0.01<br>0.45±0.01 * | 26.35±0.65<br>71.46±0.93 * | 72.49±0.64<br>28.29±0.92 * | 1.15±0.09<br>0.25±0.02 * | 1613<br>912 |

Note: Data are given as means ± standard error. * Significant differences between values at the light and dark phases.

**Supplementary 2**

**Fig. S1.** Individual periodograms of the studied endpoints at the second stage of the experiments. Significant periods on periodograms ($p<0.05$, Lomb-Scargle periodogram analysis) are above the solid red line. The red vertical line corresponds to a 27 h period. The blue vertical line corresponds to a 20 h period.

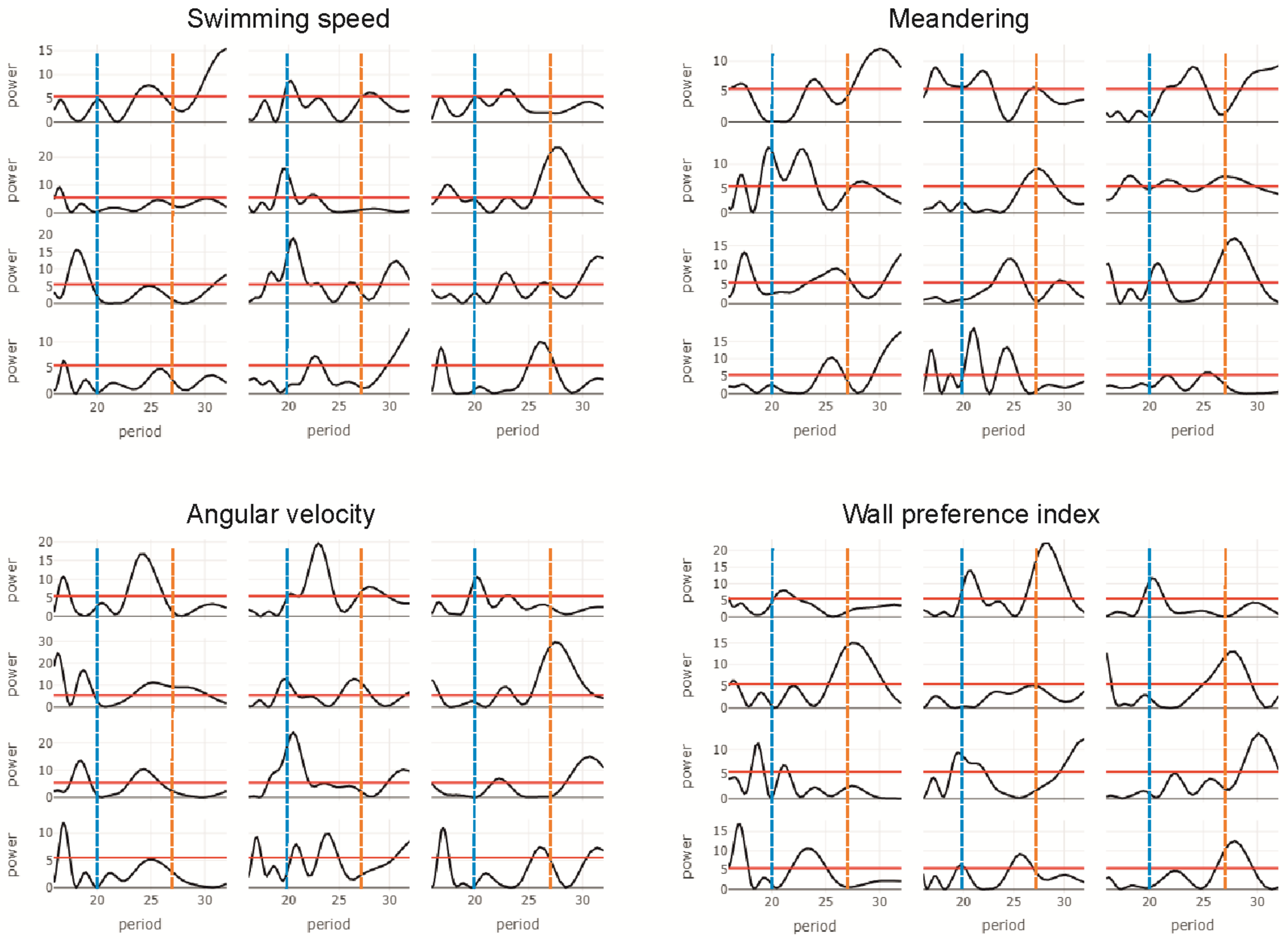

### Supplementary 3

#### Recommendations for generation of slow magnetic fluctuations

Through this study it was revealed that several conditions that must be met when reproducing the pre-recorded natural or artificially generated slow magnetic fluctuations to entrain the endogenous circadian oscillators. First of all, it is necessary to avoid a superposition of the reproduced fluctuations with natural geomagnetic variations that can affect the biological response.

1. The natural geomagnetic variation can be eliminated using the method of active compensation in the system of Helmholtz coils, as applied in this work. Alternatively, shields made of soft magnetic materials can be used. In the latter case, in addition to a zeitgeber magnetic signal, it is necessary to provide a static magnetic field corresponding to the intensity and direction of the local geomagnetic field to avoid the effects of the hypomagnetic conditions (Binhi and Prato, 2017).

Since an active compensation using feedback from a reference magnetometer can be challenging to implement, passive compensation may also be used. Though the pattern of diurnal geomagnetic variation is variable, its predictability allows for an added signal to account for these daily changes. The intensity and timing of this compensatory signal should coincide with the expected pattern of diurnal geomagnetic variation, and the direction must be opposite for each component. When reproducing the experimental signal, the diurnal variation will be compensated by fluctuations inherent in the signal. However, it should be noted that this method does not work well when a geomagnetic storm occurs or the pattern of diurnal variation changes. These events can notably modulate the resulting field affecting the test organisms. Therefore, active compensation is preferable to passive.

2. 3-component records of the diurnal geomagnetic variation during magnetically quiet times made near the experiment site can be used as the basis of signal reproduction. In this case, dynamics of natural magnetic variations, and correspondence between the artificial signal and the natural pattern of diurnal geomagnetic variation should be accounted for.

3. Sometimes (for example, when passive compensation is applied), it seems advisable to amplify the signal reproduced in the experiments to increase magnetic variation potency, separating it from possible background disturbances. The limit of natural magnetic disturbances (about 300nT) should not be exceeded as a physiological sensitivity window may exist (Wiltschko, 1978). If the standard signal were affected by sudden amplitude jumps, which may be comparable to natural geomagnetic events after amplification, test organisms may perceive these as environmental time cues. In this case, the experimental signal will need to be smoothed before amplification, leaving only the studied slow magnetic fluctuations.

4. The magnetic field dynamics during the experiments should be monitored with a magnetometer located close to the test site. It is necessary to control both the generated signals and possible temporary local disturbances of the magnetic background which can also affect the results.
